# Mechanisms for European Bat *Lyssavirus* subtype 1 persistence in non-synanthropic bats: insights from a modeling study

**DOI:** 10.1101/283564

**Authors:** Davide Colombi, Jordi Serra-Cobo, Raphaëlle Métras, Andrea Apolloni, Chiara Poletto, Marc López-Roig, Hervé Bourhy, Vittoria Colizza

## Abstract

**Background:** Lyssaviruses are pathogens of bat origin of considerable zoonotic concern being the causative agent for rabies disease, however our understanding of their persistence in bat populations remains very scarce.

**Methods:** Leveraging existing data from an extensive ecological field survey characterizing *Myotis myotis* and *Miniopterus schreibersii* bat species in the Catalonia region, we develop a data-driven spatially explicit metapopulation model to identify the mechanisms of the empirically observed persistence of European Bat Lyssavirus subtype 1 (EBLV-1), the most common lyssavirus species found in Europe. We consider different disease progressions accounting for lethal infection, immunity waning, and potential cross-species transmission when the two populations share the same refuge along the migratory path of *M. schreibersii*.

**Results:** We find that EBLV-1 persistence relies on host spatial structure through the migratory nature of *M. schreibersii* bats, on cross-species mixing with *M. myotis* population, and on a disease progression leading to survival of infected animals followed by temporary immunity. The higher fragmentation along the northern portion of the migratory path is necessary to maintain EBLV-1 sustained circulation in both species, whereas persistence would not be ensured in the single colony of *M. myotis.* Our study provides first estimates for the EBLV-1 transmission potential in *M. schreibersii* bats and average duration of immunity in the host species, yielding values compatible with previous empirical observations in *M. myotis* bats.

**Conclusions:** Habitats sharing and the strong spatial component of EBLV-1 transmission dynamics identified as key drivers in this ecological context may help understanding the observed spatial diffusion of the virus at a larger scale and across a diverse range of host species, through long-range migration and seeding of local populations. Our approach can be readily adapted to other zoonotic pathogens of public health concern.

## Background

Bats are reservoir hosts of a large number of zoonotic viral infections, including some of the recently emerged severe infectious diseases affecting humans [1–4]. They are considered to be the ancestral hosts of lyssaviruses (*Rhabdoviridae* family, *Lyssavirus* genus), the agents of rabies diseases, before the viruses in this group progressively diverged from this common ancestor to many recipient host species [5–7]. To date, bats were found to serve as reservoirs of 15 of the 17 lyssavirus species currently known [8, 9]. In Europe, four different lyssavirus species have been isolated in bats, namely *European bat lyssavirus* types 1 and 2 (EBLV-1 and EBLV-2, respectively), *Bokeloh bat lyssavirus* (BBLV), *West Caucasian bat virus* (WCBV) and one tentative species, *Lleida bat lyssavirus* [7, 10–13]. Reported in Europe for the first time in 1954 [14], EBLV-1 is the most commonly found species in the continent with a wide distribution across Germany, the Netherlands, Denmark, France, Spain [10, 14–18]. It has the potential to cross the species barrier and infect other domestic and wild mammals [14, 16, 19, 20], although such events seem relatively rare. Rabies infections of humans caused by EBLV-1 have also been reported [18], including fatal cases [13, 21].

Relatively little is known about the dynamics of EBLV-1 infection in European bats and the conditions for viral transmission and maintenance. Routine programs of passive surveillance were established in Europe at the end of the 80’s to study the distribution, abundance, and epidemiology of lyssavirus infections in bats. The vast majority of EBLV-1 positive cases was found to be associated with the Serotine bat (*Eptesicus serotinus*) [13, 14, 22], a largely diffused species in the Eurasian region that almost exclusively roost in buildings close to suburban areas, i.e. displaying a synanthropic behavior. Retrospective investigations of passive surveillance data and reports from active surveillance studies however provided evidence of EBLV-1 circulation (through neutralizing antibodies, viral RNAs) in other bat species [23], including *Pipistrellus nathusii, P. pipistrellus, Plecotus auritus*, and *Nyctalus noctula* in Germany [14], *Barbastella barbastellus, Myotis blythii, Myotis myotis, Miniopterus schreibersii* and *Rhinolophus ferrumequinum* in France [24], *E. isabellinus* (a sibling species of *E. serotinus*) in Southern Spain [25], *Rousettus aegyptiacus* in the Netherlands [26], *M. myotis* in Belgium [27], and in multiple bat species including *M. myotis* and *M. schreibersii* in Spain [25, 28, 29].

These bat species may be particularly important for the spatial diffusion and maintenance of EBLV-1 in European bats. Sequence analyses of EBLV-1 genomes from nine European countries indeed uncovered the geographic separation between phylogeographical clusters of EBLV-1 variants that cannot be fully explained by the geographic distribution of *E. serotinus* [30], a sedentary bat species [31]. Other bat species are thought therefore to be implicated in EBLV-1 circulation, with migratory species potentially assuming a prominent role in carrying the pathogen across different host populations at distant areas [32].

In addition to high mobility, the social nature of bats and their colonial aggregation constitute ideal drivers for viral exchange and dispersal [33]. Colony size and species richness were found to be associated with an increased EBLV-1 seroprevalence in Spanish bats [28, 29]. Population density and proximity of bats in the same cave provide indeed higher chances for mixing and transmission between individuals. A large number of species in the same roost might not only increase the rates of contact between bat populations, but also provide paths for cross-species diffusion and subsequent seeding events in other caves thanks to individual mobility. This suggests that infection cycles may be maintained among different host species, facilitating transient epidemics through extinction and recolonization events in a spatially structured environment, and overall potentially contributing to the spatial circulation of the virus at the regional scale.

Disease progression within individual hosts remains undefined, however it is expected to account for mechanisms to escape lethal infection and replenish the population of susceptible individuals to sustain transmission, contrasting the characteristic long lifespan of bats. Findings from passive and active surveillance suggest that bats may be capable of being exposed to lyssaviruses without dying [23]. No evidence has however addressed so far this aspect in a satisfactory or definitive way, mainly because of the scarce available knowledge on bat immunology [34]. Experimental infections in the primary host of EBLV-1 highlighted a strong dependence of the efficiency of transmission and probability of causing rabies disease on the virus transmission route [35]. The absence of virus-neutralizing activity in all sera under all transmission conditions is however in sharp contrast with field data from Spain, France, and Germany [19, 24, 28, 29, 36]. The presence of EBLV-1 neutralizing antibody response in healthy individuals appears to be relatively common and have been documented both in *E. serotinus* and in other bat species including *M. myotis* and *M. schreibersii* in various regions [24, 28, 29, 36, 37]. While its interpretation remains rather difficult, previous work suggested this response may result from bats recovering from the infection following EBLV exposure [24]. Direct evidence of transmission during abortive or subclinical infection under natural conditions is indeed difficult to achieve with active surveillance as lyssaviruses are excreted only for short periods [19, 24]. A recent longitudinal survey of *E. serotinus* colonies in France [36] found for the first time viral RNA in bats saliva concomitant with virus excretion, and later followed by seropositivity, suggesting that transmission may occur during subclinical infection. In addition, individual waves of seroconversion and waning of immunity were reported in the same colony, similar to previous results obtained for *M. myotis* in Spain [38].

Where knowledge gaps in bat ecology, epidemiology, and immunology hinder a comprehensive assessment of the mechanisms for EBLV-1 persistence in European bats, mathematical models provide a synthetic framework for hypotheses testing that can help improve our understanding of the spatial patterns reported by observational studies and identify important drivers for persistence. Here we develop a data-driven mechanistic metapopulation model for EBLV-1 spatial diffusion in the *Miniopterus schreibersii* and *Myotis myotis* non-synanthropic bat species in a system of caves in Catalonia, a region in the North-East of Spain. The model builds on data from a long-term field survey of EBLV-1 infection in natural bat colonies in the region [28, 29, 32]. *M. myotis* live as a single colony of few hundred individuals in the cave called Can Palomeres. *M. schreibersii* is a regional migratory species following a complex annual migration from cave to cave in the region. The two species share the same habitat in Can Palomeres during summer months. Through the use of spatially-resolved demographic and migration data, we explore several hypotheses regarding unknown epidemiological (transmission potential), immunological (lethal infection, immunity) and ecological aspects (cross-species mixing, seasonality in mixing, migratory behavior) to identify the mechanisms responsible for the reported EBLV-1 persistence in the two species. Given the current limitations of global surveillance for zoonotic diseases, focusing on the dynamics of bat infectious diseases and improving our understanding of the mechanisms driving their persistence may provide useful information to complement the available scarce resources to predict epizootics and potential risk for humans.

## Methods

### Metapopulation model

We develop a multi-species metapopulation epidemic model [39–41] where shelters occupied by bats are represented by model patches or subpopulations and migration events between shelters are represented by links connecting different patches. A georeferenced schematic visualization of the model is provided in the map of Figure 1. We model EBLV-1 transmission through mixing between hosts within the patches, and spatial dissemination through the migration of infected hosts. The total population includes 17,000 bats of the *M. schreibersii* species and 500 bats of the *M. myotis* species based on field estimates (see Additional File 1), and variations around these values were explored for sensitivity.

**Figure 1.**
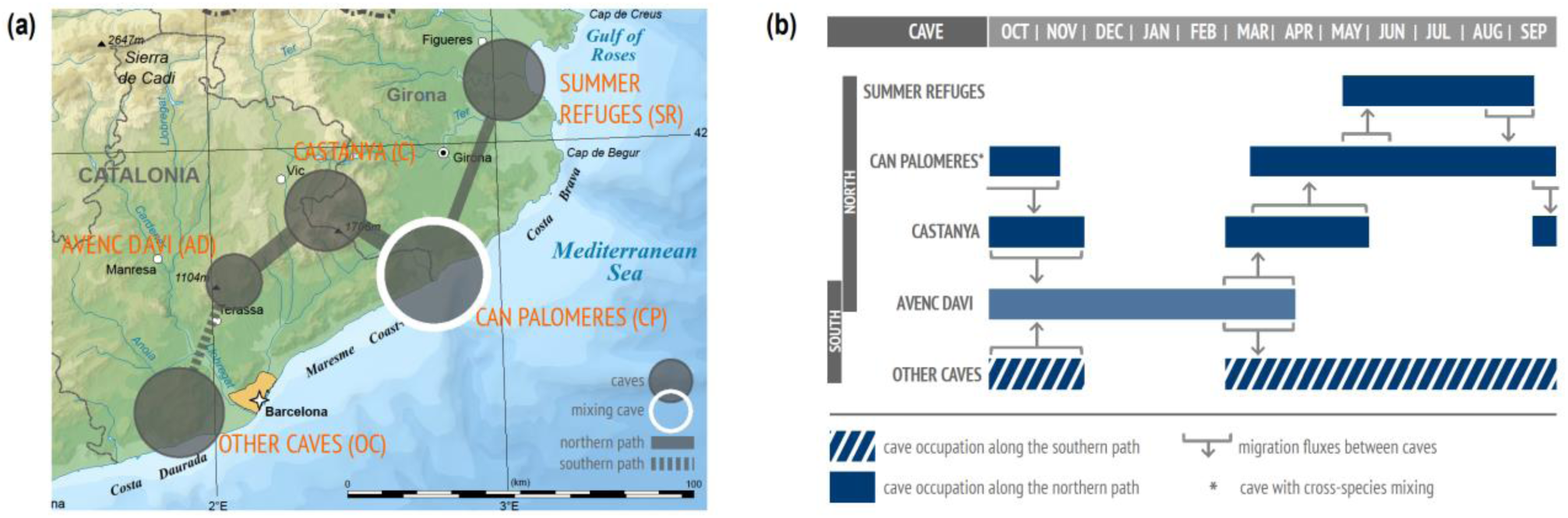
Schematic representation of the spatial model. **(a)** Georeferenced diagram of the roosting caves (circles) along the migratory path (links) of *M. schreibersii* in the region of Catalunya. Can Palomeres (white border) is the cave where cross-species mixing may occur. **(b)** Temporal representation of the annual seasonal migration of *M. schreibersii*. Cave occupation is represented with filled rectangles (northern route) and striped ones (southern route).

#### Migration

Annual migration estimates of *M. schreibersii* were obtained from a field study based on banding and recovery and presented in previous work [32]. Following hibernation in Avenc Davì (abbreviated in the following as AD), *M. schreibersii* population splits between northern and southern migration routes (Figure 1b). On the northern route, *M. schreibersii* reach Castanya (C) for mating, and then they progressively start migrating to Can Palomeres (CP) for mating and birthing. An important fraction of the population (estimated 60%) continues the migration further north to other refuges composed of breeding or summer colonies (“Summer refuges”, SR, in Figure 1a), where they stay approximately all summer. During the same period, the remaining bats stay in Can Palomeres where they share the refuge with *M. myotis*. Once summer is over, the entire *M. schreibersii* colony returns to Avenc Davì following the northern route in the opposite direction: from Summer refuges to Can Palomeres, to Castanya, to Avenc Daví for hibernation to conclude the annual migration. Bats following the southern route from Avenc Daví reach a set of caves near the coast (“Other caves”, OC) and return to Avenc Daví for hibernation, reuniting with the bats following the northern route. Migration rates are set to estimates from field data (Table S1 of Appendix File 1) [32], and a sensitivity analysis on starting date of migration events and on their duration was performed.

Since *M. myotis* bats constitute a single colony with rare and short-range movements [38], we model them as a single subpopulation staying in Can Palomeres year-round.

Consistently with the cycle described above, we consider a seasonal birth pulse during the birthing summer season in Can Palomeres for both species, and constant natural death rate.

#### EBLV-1 infection dynamics

We propose three models for EBLV-1 infection dynamics in the bat species under study, to account for different hypotheses based on field observations and experimental knowledge. They are all based on a susceptible-exposed-infected-recovered compartmental scheme [42], with variations to account for different immunological responses. Model 1 assumes non-lethal infection and loss of immunity (panel a of Figure 2), as done in previous modeling works [38, 43]. Model 2 considers the possibility for bats to develop a lethal infection (with a given probability *ρ*), alternative to a non-infectious state followed by permanent immunity (panel b). Model 3 is a variation of model 2 that considers temporary immunity (panel c).

**Figure 2.**
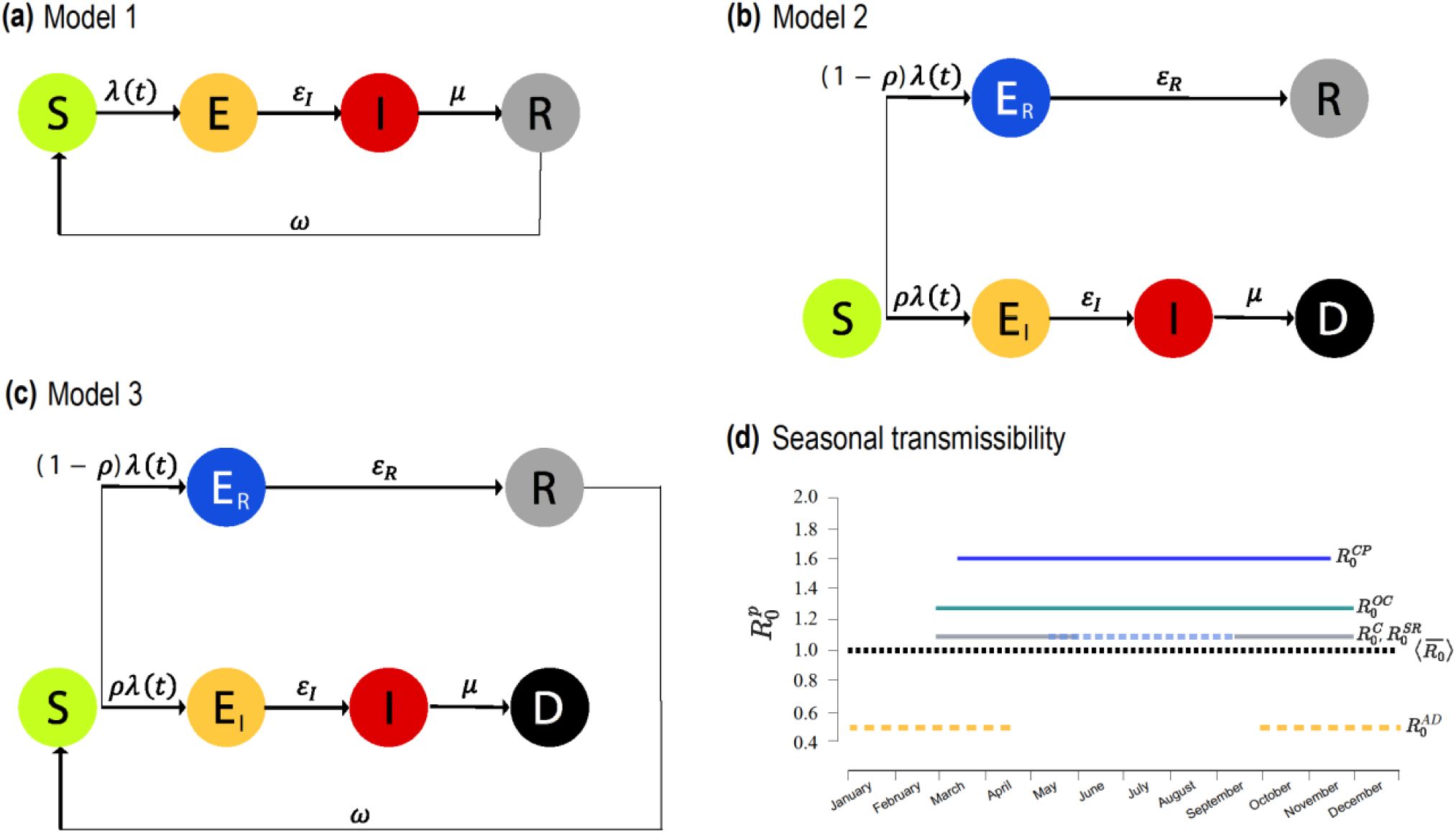
Disease progression models and seasonality of transmission. **(a)** Compartmental structure for model 1, where no infection-induced mortality is considered and immunity wanes with rate *ω*. *ε*_*I*_ is the rate of becoming infective following infection, and *μ* the recovery rate **(b)** Compartmental structure for model 2, considering lethal infection to occur with probability *ρ*, whereas non-lethally exposed individuals (*E*_*R*_) recover with rate *ε*_*R*_ to the permanently immune state. **(c)** As in (b) for model 3, where immunity wanes with rate *ω*. Demographic processes in the three diagrams are omitted for clarity. **(d)** Reproductive numbers 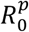 for *M. schreibersii* along each patch *p* of the migration path. The values correspond to the maximum likelihood estimates. The average reproductive number of the metapopulation model, 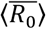, is also shown (black dashed curve).

Seasonality affects transmission intensity, as it varies upon the degree of bats activity throughout the year. For *M. schreibersii*, we model it through a patch-dependent variation of the reproductive number *R*_0_, a key epidemiological parameter measuring the average number of secondary cases that an infectious host can generate during the infectious period in a fully susceptible population [44]. We consider the reproductive number 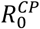 in Can Palomeres as an input to the model and corresponding to the highest transmissibility, associated with the highest mixing for mating and birthing occurring in that cave. Reproductive numbers in the other patches assume smaller values: 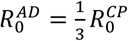 in Avenc Davì (i.e. the smallest value during hibernation period) and 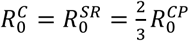 in Castanya and Summer refuges, based on expert opinion. For Other caves along the southern route we make the parsimonious hypothesis of 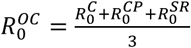, equal to the average value of northern’s path parameters. The modeled seasonal variation of transmission intensity in each cave for *M. schreibersii* is reported in Figure 2d. Robustness of our findings against these assumptions was tested through the comparison with an experimental scenario lacking seasonality (see *Experimental scenarios* subsection). For *M. myotis*, we model seasonality in transmission intensity as a two-step function describing hibernation in winter months (as for *M. schreibersii* in Avenc Davì) and breeding and mating during the rest of the year (as for *M. schreibersii* in Can Palomeres).

Cross-species transmission between *M. schreibersii* and *M. myotis* may occur in Can Palomeres only. We model it with a reduction in the transmissibility, 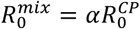 with 0 ≤ *α* ≤ 1, to account for reduced mixing between different species; *α* = 0 refers to non-mixing conditions. As an example, we provide here the force of infection for *M. schreibersii* in Can Palomeres in model 1, whereas full details for all models are reported in the Additional File 1:

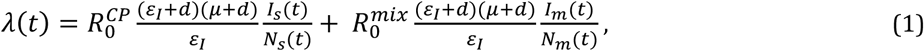

where *ε*_*I*_ is the rate of becoming infective following infection, *μ* the recovery rate, *d* the mortality rate, *I*_*s*_(*t*) is the number of infective *M. schreibersii* bats in the Can Palomeres population of size *N*_*s*_(*t*) at time *t*, and analogous (*I*_*m*_(*t*), *N*_*m*_(*t*)) for *M. myotis*.

We set disease progression parameters for *M. myotis* to available estimates from previous studies [38, 43]. Lacking data characterizing the disease progression of EBLV-1 in *M. schreibersii*, we set the average durations of the incubation and infectious period to the values estimated for *M. myotis*, following previous modeling work [43]. Variations of these values are then considered for sensitivity analysis. We explore three values of the probability *ρ* of lethal infection (for models 2 and 3), *ρ* = 0.15, 0.35, 0.5, to consider limited, moderate, and relatively high probabilities, in absence of estimates from field data. The reproductive number 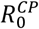 (for all models) and the average immunity period *ω*^−1^ (for models 1 and 3) are unknown but expected to play a crucial role in determining EBLV-1 persistence in the two host populations, therefore we explore the following ranges of values: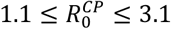, and 180 ≤ *ω*^−1^ ≤ 780 days.

Parameters with their description and values are reported in Table S5 in the Additional File 1.

### Numerical simulations and persistence analysis

We perform discrete stochastic numerical simulations of the EBLV-1 transmission in the host populations to account for the discrete nature of hosts and for stochastic extinction events that may be favored by small host population sizes. Time is considered to be discrete with a daily timescale. The epidemic is seeded with 100 infected *M. schreibersii* bats and 10 infected *M. myotis* bats in the hibernation period, and different initial conditions are explored for sensitivity. For each model and under each hypothesis considered, we ran 10^3^ stochastic simulations starting from the same initial conditions and reaching the endemic equilibrium. Simulations provide at each time step the number of *M. schreibersii* and *M. myotis* in each compartment in each cave, and the number of *M. schreibersii* migrating from one cave to another. The metapopulation framework is implemented in C++, and technical details for simulations are reported in the Additional File 1.

We compute the persistence probability of EBLV-1 in each bat species as the fraction of stochastic simulations reaching the endemic equilibrium in both host populations. To compare numerical results across different models and hypotheses, we consider a metapopulation summary measure for *M. schreibersii* species given by the average reproductive number of the metapopulation model across time and caves:

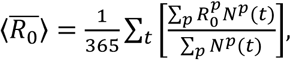

where 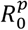 represents the reproductive number of patch *p*, and *N*^*p*^(*t*) indicates the *M. schreibersii* population size of patch *p* at time *t*.

We use a maximum likelihood approach to compare the seroprevalence data from the two species [45] with our numerical results to identify values and associated confidence intervals of the reproductive number 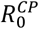 in Can Palomeres and of the immunity period for *M. schreibersii* mostly compatible with observations. Details are reported in the Additional File 1.

### Experimental scenarios

To assess the impact of several ecological drivers on the probability of persistence of EBLV-1 in both species, we compare our data-driven metapopulation model with a set of experimental scenarios that we describe here.

To evaluate the role of seasonality in transmission, we build a *no-seasonal metapopulation* epidemic model with the same spatial structure of the data-driven metapopulation model (i.e. patches, demographics and migration dynamics of Figure 1), but with no variation in the transmissibility associated to the caves. The reproductive number is assumed to be constant in time and space, and equal to 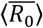.

To identify the portions of the migration path that are mostly relevant for disease persistence, we consider a *northern path only* metapopulation model considering only bats following the path including Avenc Davi, Castanya, Can Palomeres and Summer refuges, and a *southern path only* metapopulation model considering instead only bats following the path from Avenc Davi to Other caves and return. Time and rates of migration events remain as in the full migration path.

To assess the role of spatial resolution in the identification of patches and associated mixing opportunities, we consider a *higher resolution* metapopulation model where all caves in the Summer refuges are independently considered as patches (corresponding migration flow estimates are provided in the Additional File 1).

Finally, we test density-dependent transmission rates for EBLV-1 dynamics in both species, alternative to the frequency-dependent assumption considered in Eq. (1), to explore different mixing conditions.

## Results

EBLV-1 circulation in both species is numerically recovered only in model 1 (temporary immunity and non-lethal infection) with cross-species mixing (Figure 3, panels a and b). The absence of mixing between *M. schreibersii* and *M. myotis* or lethal infection lead instead to negligible or null probability of persistence in one of the species (*M. myotis*, Figure 3, panel d) or both (Table S6 of the Additional File 1, for different probabilities leading to the lethal infectious state), respectively. Also, density-dependent transmission would not allow persistence of the pathogen in any of the models explored (Table S6).

**Figure 3.**
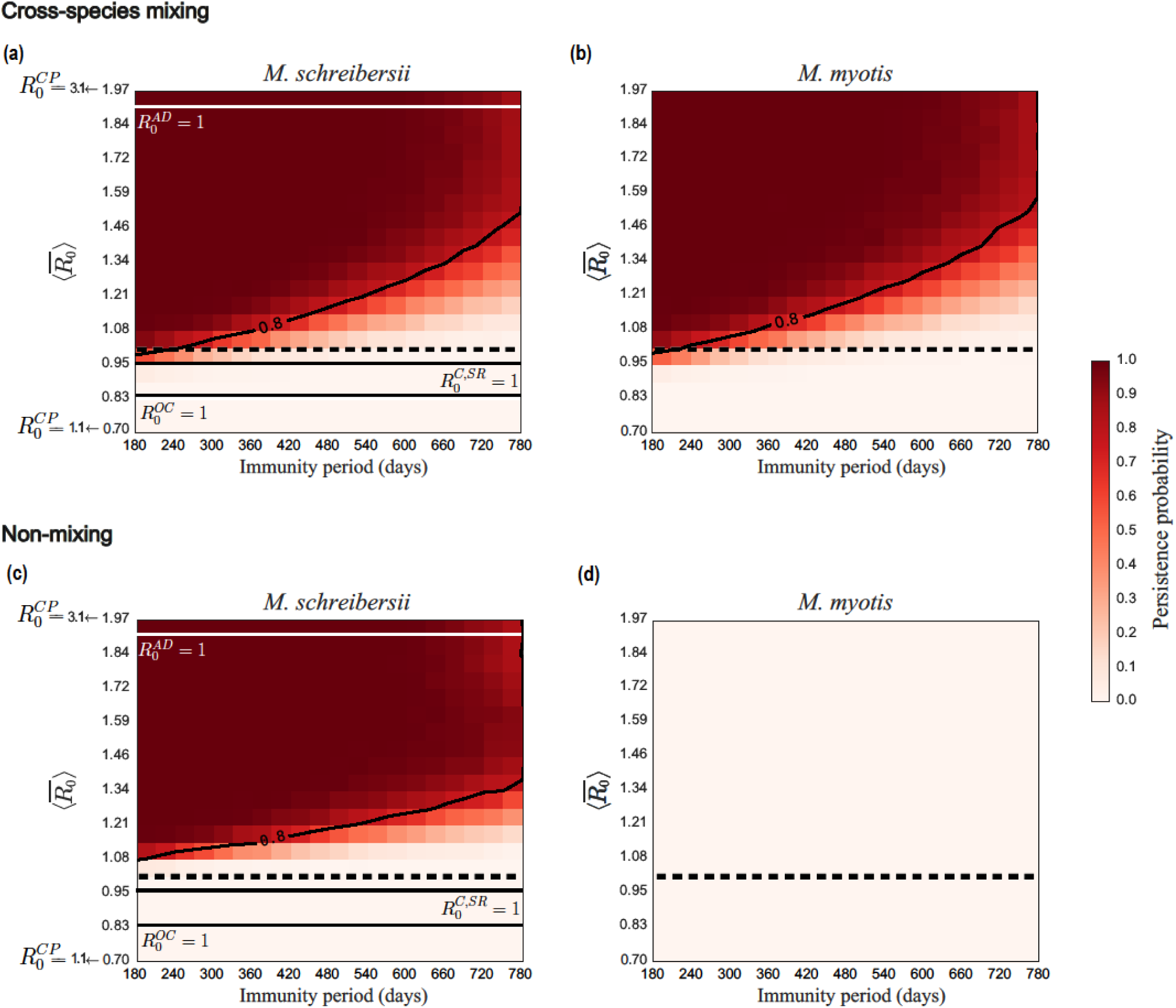
Persistence probability of EBLV-1 in *M. schreibersii* and in *M. myotis* bats in model 1. (a), (b): Persistence probability for *M. schreibersii* (a) and for *M. myotis* (b) as a function of the average reproductive number of the metapopulation model 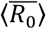 and of the immunity period *ω*^−1^ in the mixing scenario. **(c), (d):** as in (a), (b) in the non-mixing conditions. Contour lines indicate a persistence probability of 80%. The dashed horizontal line refers to 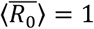. Solid horizontal lines refer to threshold conditions 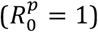 for the caves.

Persistence probability profiles are very similar in the two host populations in model 1 with cross-species mixing. Virus circulation is maintained for all values of the immunity periods explored, with high persistence ensured at lower transmissibilities if the immunity period is short. Mixing allows persistence also for values of the average metapopulation reproductive number close to or below the critical threshold, favoring viral circulation compared to the non-mixing case (Figure 3, panels a and c).

Comparison with serological data is conducted for model 1 with cross-species mixing, i.e. the sole modeling framework predicting persistence in both species, as observed in nature. Best estimate values for the unknown parameters are *ω*^−1^ = 390 (95% CI: 228-772) days and 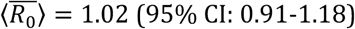 (95% CI: 1.43-1.84). The latter is associated to a metapopulation average just above the critical threshold, 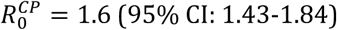, and the reproductive number is predicted to be largely subcritical in Avenc Davi during the hibernation period, 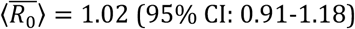 (Table 1). With 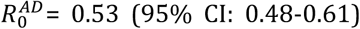 set to its maximum likelihood estimate, we find the probability of viral persistence in *M. schreibersii* to vary strongly, decreasing from 82% to 1% when the immunity period varies within its 95% confidence interval, with a probability equal to 38% for the best estimate *ω*^−1^ = 390 days (Figure S1 in the Additional File 1). Such trend in the persistence probability is not substantially altered by variations in the assumed values for cross-species mixing (Figure S2 in the Additional File 1).

**Table 1.**
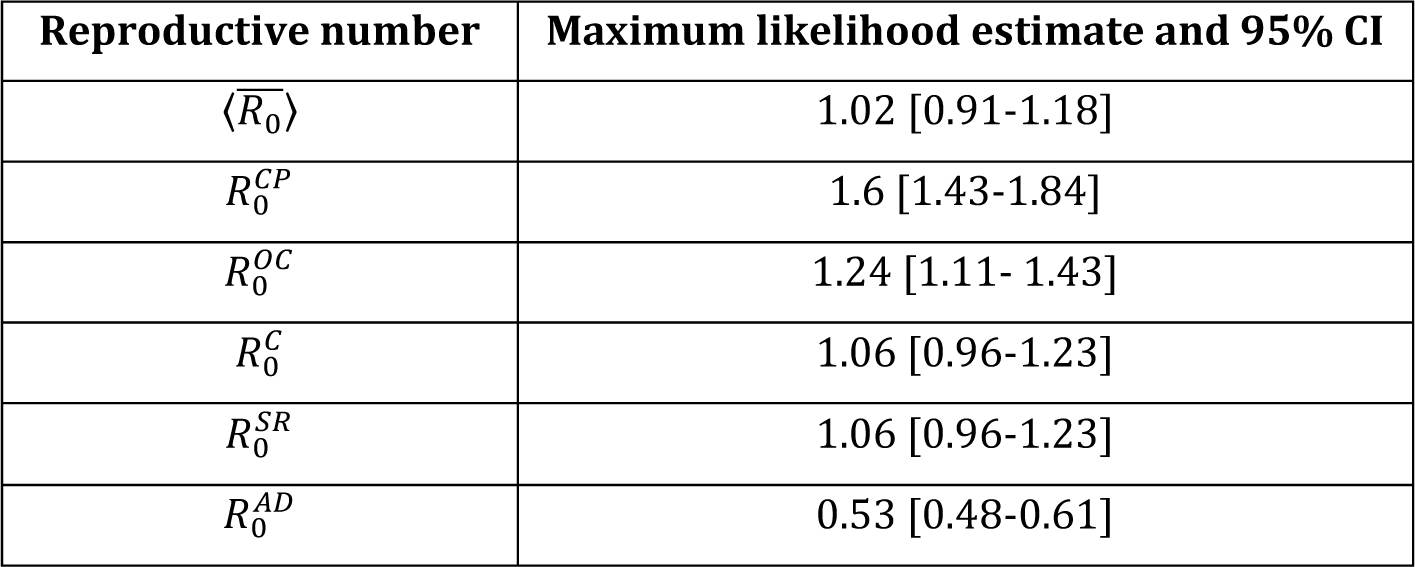
Maximum likelihood estimates for the reproductive number.

| Reproductive number | Maximum likelihood estimate and 95% CI |
| --- | --- |
| $\langle \bar{R}_0 \rangle$ | 1.02 [0.91-1.18] |
| $R_0^{CP}$ | 1.6 [1.43-1.84] |
| $R_0^{OC}$ | 1.24 [1.11- 1.43] |
| $R_0^C$ | 1.06 [0.96-1.23] |
| $R_0^{SR}$ | 1.06 [0.96-1.23] |
| $R_0^{AD}$ | 0.53 [0.48-0.61] |

The analysis of the experimental scenarios allows us to identify the ecological drivers that are important for persistence (Figure 4). Discarding yearly seasonality of transmission leaves the persistence probability almost unaltered. The essential role of migration is ensured by its northern portion, without which the likelihood of viral maintenance would be strongly reduced for a large set of values of 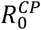, becoming null when 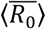 assumes its maximum likelihood estimate. Finally, considering the breakdown of the Summer refuges patch into smaller subpopulations, i.e. through a higher spatial resolution metapopulation model, would require a slight increase in transmissibility to reach the same persistence values of the reference model.

**Figure 4:**
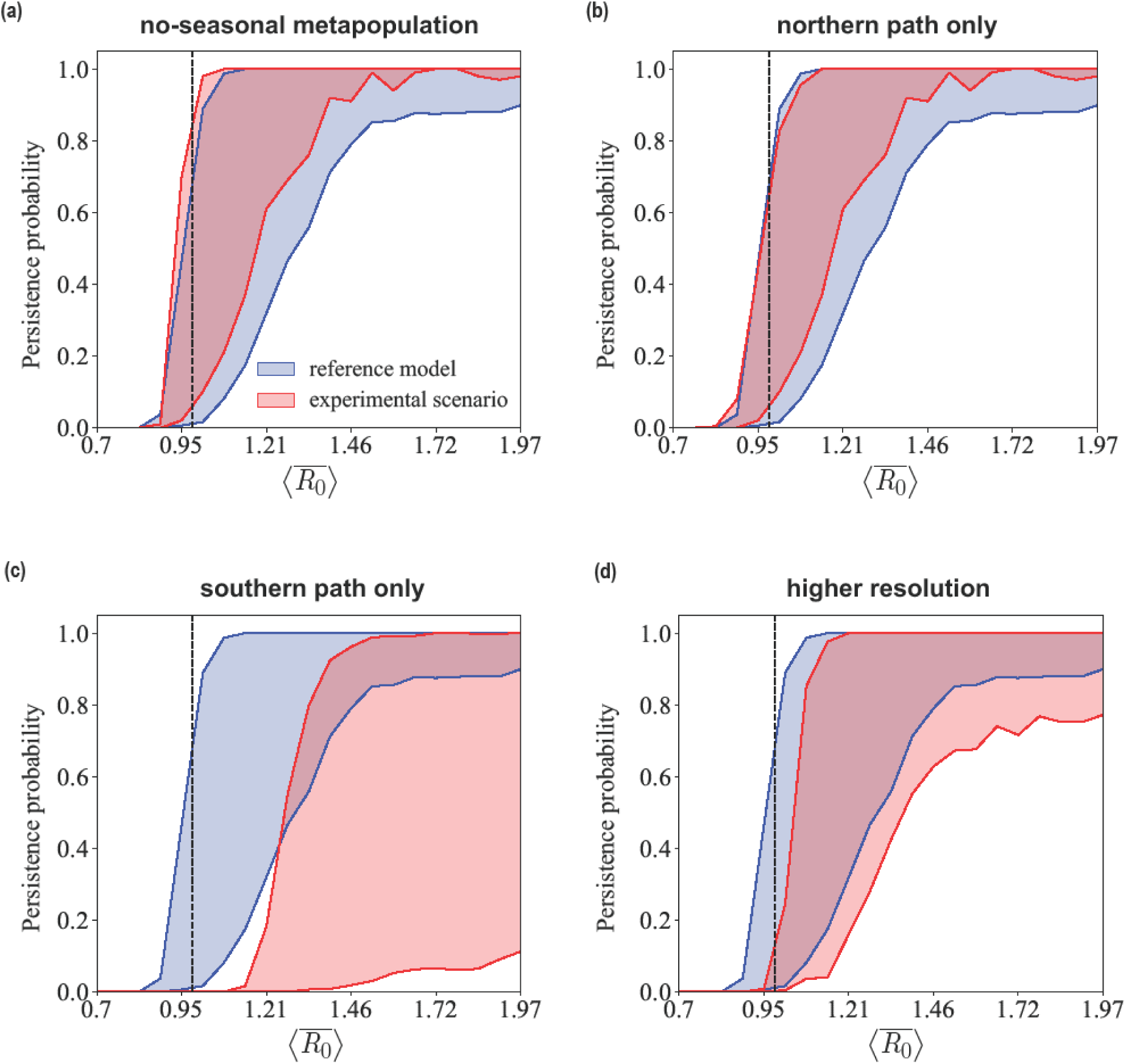
Comparison with experimental scenarios. Persistence probability for *M. schreibersii* as a function of the average reproductive number of the metapopulation model 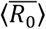 for values of the immunity period *ω*^−1^ spanning the estimated confidence interval. Each experimental scenario indicated in the plot title is compared with the reference model, corresponding to the data-driven metapopulation model. Numerical results are obtained for model 1 in mixing conditions.

We performed a sensitivity analysis to allow for variations in the ecological estimates (bat population sizes, Figures S4 and S5 of the Additional File 1; starting date of migration events, Figures S6; duration of migration events, Figures S7), and in the assumed length of the infectious period (Figures S3) yielding no variation in the predicted conditions for EBLV-1 circulation in both species.

## Discussion

Through a spatially explicit multispecies metapopulation model based on data from a long-term field survey on *M. schreibersii* and *M. myotis* bats in Spanish natural colonies, we were able to identify the main drivers for EBLV-1 persistence in the ecosystem under study and provide novel numerical evidence informing on previously unknown epidemiological, immunological and ecological factors.

Overall our findings indicate that EBLV-1 persistence relies on host spatial structure through the migration of *M. schreibersii* bats, on cross-species mixing with *M. myotis* population, and on a disease progression leading to survival of infected animals followed by temporary immunity. Even a low probability of developing lethal infection together with bats migration and reseeding through cross-species mixing would not be able to sustain the epidemic, regardless of the loss of immunity. While Lyssavirus infections are known to be generally lethal for mammals and for some bat species [46, 47], current knowledge from experimental and natural studies is not sufficient to accurately and satisfactorily define EBLV-1 disease progression in bats in natural conditions [24, 28, 29, 34, 36, 37]. This is further complicated by biological and regulatory aspects. First, despite recent evidence for *E. serotinus* bats in France [36], detecting a subclinical infective state in healthy bats in natural colonies is rather unlikely because of the short duration of the excretion period, and no such evidence exists yet for *M. schreibersii* or *M. myotis.* Second, the legal framework protecting European bats [48] makes field studies particularly difficult to implement, as for instance marking bats is forbidden in Europe and special authorizations need to be requested to conduct these studies. Little but precious available field data are however providing an increasing body of evidence pointing to the possibility that various bat species may experience an infective subclinical state after being exposed to EBLV-1, followed by recovery and loss of immunity, with no associated increased mortality [19, 24, 29, 36, 38], in agreement with our model results. A constant survival rate despite recurrent EBLV-1 epidemic cycles was reported for *M. myotis* [38], supporting the results of our model. Additional field data is needed to confirm our numerical predictions on EBLV-1 infection in *M. schreibersii.*

Two ecological factors emerge as critical drivers for viral persistence: cross-species mixing and host migration. We found that EBLV-1 circulation would not be maintained in a single colony alone in absence of mixing with other species. Multi-species colonies are indeed a phenomenon largely observed in the field that is known to favor virus exchange [28, 49]. The importance of cross-species transmission of lyssaviruses was also reported in other ecological settings, identifying phylogenetic distance as the key determinant for cross-species transmission of rabies virus in North American bat species [5, 50]. No study of this kind has been performed yet on EBLV-1 hosts species, and while the mixing intensity between *M. schreibersii* and *M. myotis* in Can Palomeres is unknown, our persistence estimates remain quite robust against this assumption. Our findings suggest that an increased public health attention should be focused on caves hosting multiple bat species with targeted ecological fieldwork to improve our understanding of the role of species richness in viral exchange.

We find that transmission may not depend on host density in the cave. Many bat species are indeed known to form communities (families) that are stable over short terms (daily and nocturnal activities) and long terms (between migrations) thus including members of different generations, and whose sizes are independent on the colony size [51, 52]. Given that virus transmission through bites and scratches [35] requires close contact, the regularity of interactions with a limited number of hosts may explain the frequency-dependent transmission selected by our model. This result however seems to depend considerably on the species considered and the roosting behavior [53]. Additional modeling work focusing on different contexts might help better understanding such dependence across different conditions.

The other component critical for stable EBLV-1 circulation is the presence of the migratory species. Movements of infected individuals provide a mechanism for maintaining the chains of transmission through the seeding of epidemics in different patches. The importance of frequent immigration of infected hosts for persistence was also recognized in other settings [43, 45, 54, 55]. Moreover, our findings indicate that the migratory species may contribute to pathogen persistence in species encountered along the migratory path. Given the potential for long-range seasonal movements of *M. schreibersii* [29, 32], this species may represent a central vector for spatial dispersion of EBLV-1 in southern Europe, where this bat species is abundant, possibly contributing to reconcile the discrepancy observed between the phylogeographical clusters of EBLV-1 variants and the geographic distribution of its common host species *E. serotinus* [30].

In the ecological context under study, the northern portion of the migratory path is entirely responsible for viral persistence at the estimated immunity waning, likely because it is composed by a more complex spatial structure including a larger number of patches, thus creating more opportunities for seeding events sustaining coupled but not synchronous patch epidemics [56] that cannot be otherwise obtained with the southern path only. Increasing spatial resolution and resolving shelters sharing similar ecological and environmental conditions (such as the ones collectively grouped in Summer refuges) would not substantially alter our predictions. These findings are important to inform future field studies minimizing data collection efforts on roosts occupation.

Seasonality in mixing between hosts was instead found to have a negligible impact on EBLV-1 maintenance, suggesting that field efforts should be prioritized to provide an accurate characterization of the migration pattern, an important driver for EBLV-1 endemic circulation, instead of hosts’ degree of interaction.

The maximum likelihood analysis allows us to provide for the first time previously unidentified parameters characterizing the disease dynamics of EBLV-1 in *M. schreibersii.* We find that transmission among *M. schreibersii* bats in Can Palomeres (i.e. under the highest mixing conditions) is similar to what was previously estimated for *M. myotis* (*R*_0_ = 1.7) in natural colonies in Spain [38]. These conditions of transmissibility are associated to other caves being in close-to-critical or sub-critical conditions for efficient epidemic transmission. Persistence is thus mainly supported by transmission in Can Palomeres, though this cave only is not sufficient to maintain endemicity in *M. myotis*. The modeling predictions for the immunity period are in the ballpark of previous empirical estimates of the maximum length of seropositive status observed in *M. myotis* (2-3 years) [29] and in *E. serotinus* (4 years) [36]. Individuals are predicted to be immune on average for more than a year, thus hindering virus survival because of the slow replenishment of susceptible hosts. This effect is however counterbalanced by the migration of hosts that occurs on an annual timescale, providing opportunities for seeding events in naïve populations, a mechanism already identified by theoretical works to sustain viral circulation [56]. A rather large confidence interval is obtained for the estimate of the immunity period, indicating that the available serological data are not sufficient to significantly discriminate between approximately 1 year and 2 years of duration of immunity. Also, the likelihood of persistence is predicted to vary quite considerably within this range, as the system is found in the transition between null or negligible persistence and very high probability of viral maintenance. Additional cross-sectional studies in this colony and at higher temporal resolution may help improve our estimates.

The robustness of our model to a set of ecological, epidemiological, and immunological assumptions highlights that, despite the very limited knowledge of the system, our data-driven approach is able to clearly identify the mechanisms underpinning EBLV-1 circulation in the studied populations, providing important avenues for further empirical investigations.

Our study is affected by some limitations. We did not consider *E. serotinus* bats in our model, the host species associated with the large majority of EBLV-1 cases detected in Europe [14, 22, 57]. While present in the region under study, its synanthropic behavior (i.e. living close to humans populations) likely precluded interactions with *M. schreibersii* and *M. myotis*, which instead clearly exhibit a non-synanthropic behaviour, roosting in natural caves and in abandoned mines [29]. Moreover, our model considers the two host populations to be closed and isolated in the region under study. While contacts with other non-synanthropic bat species of smaller population sizes may occur in Can Palomeres, our findings show that 2 host species are enough to self-sustain the epidemic given cross-species mixing and 1-species migration, with no need for additional introductions from other sites. This may appear to be in contrast with was found in other contexts, e.g. in a rabies virus model in vampire bats where persistence was largely dependent on immigration of infected individuals [45]. Our model however already accounts for seeding events from one patch to another thanks to the migration of *M. schreibersii*. Similarly to the study by Blackwood et al., we find indeed that the virus would not be maintained in a single population only.

## Conclusions

Bat species have a wildly variable range of habitats, life cycles, population sizes and spatial distributions. Some species display a more localized nature like *M. myotis*, while others exhibit rather long migratory behaviors similar to *M. schreibersii.* Based on data from a long-term field survey, our work identifies cross-species mixing and host migration as determinants for EBLV-1 circulation in these two populations, as long as infection with the virus is not lethal for the animals. The uncovered strong spatial component to transmission dynamics may help understanding the observed spatial diffusion of the virus at a larger scale and across a diverse range of host species, through long-range migration and seeding of local populations. Though the risk of lyssavirus infection to humans remains relatively low, the invariably fatal outcome of clinical disease in humans makes it an important issue for public health. Our framework can readily be extended also to other zoonotic viral pathogens of public health concern circulating in spatially dispersed bat populations.

## Abbreviations

EBLV-1: European bat lyssavirus subtype 1
EBLV-2: European bat lyssavirus subtype 2
BBLV: Bokeloh bat lyssavirus
WCBV: West Caucasican bat virus
AD: Avenc Davì
C: Castanya
CP: Can Palomeres
SR: Summer refuges
OC: Other caves
CI: Confidence interval

## Declarations

### Ethics approval and consent to participate

Not applicable

### Consent for publication

Not applicable

### Availability of data and materials

Additional file 1. provides empirical data, mathematical formulation of the models, and additional numerical results..

### Authors’ contributions

Conceived the experiments: HB, JSC, AA, CP, VC. Designed the experiments: DC, RM, VC. Performed the experiments: DC, RM, AA. Analyzed the data: DC, RM, AA, MLR, JSC, HB. Wrote the manuscript: DC, RM, VC. All authors gave final approval for publication.

## Acknowledgements

We thank Lulla Opatowski and Pierre-Yves Böelle for useful discussions.

