## Supplementary Materials for "Mechanisms for European Bat *Lyssavirus* subtype 1 persistence in non-synanthropic bats: insights from a modeling study"

**ADDITIONAL FILE 1**

### 1. Empirical data

Empirical data on demographic sizes of *M. schreibersii* and *M. myotis* colonies (Table S1) and on migration movements of *M. schreibersii* (Table S2) were obtained from a long-term survey of bat populations in natural colonies in Spain. The data we present hereafter are a continuation of the fieldwork and results already presented in Refs. [1, 2], so we refer the reader to those papers for details on data collection.

**Table S1. Bat population estimates.**

| Sampling date | *Miniopterus schreibersii* population estimate in Avenc Davi | *Myotis myotis* population estimate in Can Palomeres |
|---|---|---|
| Dec 2009 | 18,050 | 537 |
| Dec 2010 | 17,100 | 577 |
| Dec 2011 | 17,100 | 460 |
| Dec 2012 | 17,100 | 441 |
| Dec 2013 | 15,200 | 528 |
| Dec 2014 | 17,100 | 520 |
| Dec 2015 | 18,050 | 565 |
| Dec 2016 | 17,100 | 575 |
| Average and 95% CI | 17,100 [16,490 – 17,709] | 525 [489 - 560] |

**Table S2. Migration estimates for *Miniopterus schreibersii.***

| Origin cave | Destination cave | % migrating bats | Start date | Duration (days) | Daily migration rate |
|---|---|---|---|---|---|
| Avenc Davì | Castanya | 64 | 1st March | 46 | 0.0217 |
| Avenc Davì | Other caves | 36 | 1st March | 46 | 0.0217 |
| Castanya | Can Palomeres | 100 | 15th March | 78 | 0.0128 |
| Can Palomeres | Summer refuges | 60 | 15th May | 31 | 0.0322 |
| Summer refuges | Can Palomeres | 100 | 10th August | 36 | 0.0277 |
| Can Palomeres | Castanya | 100 | 15th September | 61 | 0.0164 |
| Castanya | Avenc Davì | 100 | 1st October | 60 | 0.0166 |
| Other caves | Avenc Davì | 100 | 1st October | 60 | 0.0166 |

In the higher resolution experimental scenario, we model explicitly the migration within different caves denoted as Summer refuges. In the following Table we provide the estimate for the associated migration flows.

**Table S3. Migrations estimates for *Miniopterus schreibersii* within Summer refuges.**

| Origin cave | Destination cave | % migrating bats | Start date | Duration (days) | Daily migration rate |
|---|---|---|---|---|---|
| Can Palomeres | Frare | 70 | 15th May | 31 | 0.0322 |
| Can Palomeres | Ratapenades | 30 | 15th May | 31 | 0.0322 |

EBLV-1 serological data were collected once a year, in Avenc Davì (for *M. schreibersii*) and Can Palomeres (for both species), between 2010 and 2014 (Table S3). From the 279 *M. schreibersii* and 25 *M. myotis* sampled, 15.41% (95% CI [11.49-20.31], n=43) and 40% (95% CI [21.81-61.11], n=10) were EBLV-1 seropositive, respectively. These analyses represent the continuation of previous fieldwork, so we refer the reader to Refs. [2–4] for details on bat sampling and detection of EBLV-1 neutralizing antibodies.

While estimates are yielded from relatively small sample sizes, they are rather consistent across the 5 years of the study and with the estimates obtained from previous portions of the long-term survey [2–4]. Positive animals were found every year, except in 2014 when only one animal of the *M. myotis* species was sampled. Also, sampling rates are compatible with other field surveys in Europe [5–7].

**Table S4. Number of bats sampled and tested positive to EBLV-1 serology, prevalence and 95%CI, in Avenc Davi and Can Palomeres, for both bats species.**

| Sampling date | Cave | Species | Sampled | Positive | Prevalence [95%CI] |
|---|---|---|---|---|---|
| 23/03/10 | Avenc Davì | *M. schreibersii* | 29 | 3 | 0.10 [0.04-0.26] |
| 24/03/11 | Avenc Davì | *M. schreibersii* | 31 | 4 | 0.13 [0.05-0.29] |
| 28/03/12 | Avenc Davì | *M. schreibersii* | 30 | 3 | 0.10 [0.03-0.26] |
| 28/03/13 | Avenc Davì | *M. schreibersii* | 31 | 0 | 0.00 [0.00-0.11] |
| 20/03/14 | Avenc Davì | *M. schreibersii* | 29 | 0 | 0.00 [0.00-0.12] |
| 22/07/10 | Can Palomeres | *M. schreibersii* | 24 | 13 | 0.54 [0.35-0.72] |
| 09/06/11 | Can Palomeres | *M. schreibersii* | 26 | 0 | 0.00[0.00-0.13] |
| 16/05/12 | Can Palomeres | *M. schreibersii* | 30 | 14 | 0.47 [0.30-0.64] |
| 03/07/13 | Can Palomeres | *M. schreibersii* | 22 | 2 | 0.09 [0.03-0.28] |
| 05/06/14 | Can Palomeres | *M. schreibersii* | 27 | 4 | 0.15 [0.06-0.32] |
| 22/07/10 | Can Palomeres | *M. myotis* | 6 | 3 | 0.50 [0.19-0.81] |
| 09/06/11 | Can Palomeres | *M. myotis* | 4 | 1 | 0.25 [0.01-0.70] |
| 16/05/12 | Can Palomeres | *M. myotis* | 5 | 3 | 0.60 [0.23-0.88] |
| 03/07/13 | Can Palomeres | *M. myotis* | 9 | 3 | 0.33 [0.12-0.65] |
| 05/06/14 | Can Palomeres | *M. myotis* | 1 | 0 | 0.00 [0.00-0.95] |

### 2. Mathematical formulation of the models and parameter values

#### 2.1 Model 1

Model 1 assumes the following transmission dynamics: infected bats become exposed ($E$), then infectious ($I$) and able to transmit the disease, and recover with rate $\mu$. After an average immunity period $\omega^{-1}$, they become again susceptible. Disease progression is mathematically described by the following equations, written in deterministic continuous differential equations for sake of simplicity. Simulations are instead discrete and stochastic (see Section 3).

$$\frac{dS^p}{dt} = -\beta^p \frac{S^p I^p}{N^p} - dS^p + \omega R^p$$

$$\frac{dE^p}{dt} = \beta^p \frac{S^p I^p}{N^p} - (\varepsilon_I + d)E^p$$

$$\frac{dI^p}{dt} = \varepsilon_I E^p - (\mu + d)I^p$$

$$\frac{dR^p}{dt} = \mu I^p - (\omega + d)R^p$$

where $N$ is the total bats population and the superscript $p$ refers to the patches. In Can Palomeres we model the birth rate pulse $b$, and we add the cross species interaction through the factor $\alpha\beta^{CP}S^{CP}I_{s2}^{CP}$, where $I_{s2}^{CP}$ indicates the number of infected of the other species and $\alpha$ is the cross-species mixing interaction:

$$\frac{dS^{CP}}{dt} = bN^{CP} - \beta^{CP}S^{CP}\left(\frac{I^{CP}}{N^{CP}} + \alpha\frac{I_{s2}^{CP}}{N_{s2}^{CP}}\right) - dS^{CP} + \omega R^{CP}$$

$$\frac{dE^{CP}}{dt} = \beta^{CP}S^{CP}\left(\frac{I^{CP}}{N^{CP}} + \alpha\frac{I_{s2}^{CP}}{N_{s2}^{CP}}\right) - (\varepsilon_I + d)E^{CP}$$

$$\frac{dI^{CP}}{dt} = \varepsilon_I E^{CP} - (\mu + d)I^{CP}$$

$$\frac{dR^{CP}}{dt} = \mu E^{CP} - (\omega + d)R^{CP}$$

The intrinsic $R_0^p$ value, the basic reproductive number in the absence of migration per each patch $p$, is obtained using the next generation matrix approach [8]:

$$R_0^p = \frac{\beta^p\varepsilon_I}{(\varepsilon_I + d)(\mu + d)} \tag{1}$$

The force of infection for intra-species infection in a given patch $p$, at time $t$ is:

$$\lambda^p(t) = R_0^p\frac{(\varepsilon_I + d)(\mu + d)}{\varepsilon_I}\frac{I^p(t)}{N^p(t)} \tag{2}$$

In Can Palomeres, where cross-species transmission between *M. schreibersii* and *M. myotis* leads to an additional contribution to the force of infection, $R_0^{mix}\frac{(\varepsilon_I + d)(\mu + d)}{\varepsilon_I}\frac{I_{s2}^{CP}(t)}{N_{s2}^{CP}(t)}$, where $R_0^{mix} = \alpha R_0^{CP}$. In our numerical simulations, we fix a value for $R_0^{CP}$ (and then explore a full range of values), compute the associated reproductive numbers $R_0^p$ in all other patches, and calculate the corresponding forces of infection.

### 2.2 Model 2

Model 2 assumes the following transmission dynamics: once infected, bats can experience a lethal infection with rate $\rho$, becoming exposed ($E_I$), then infectious ($I$) and finally dying with rate $\mu$. With a non-lethal infection (probability $(1-\rho)$), they become exposed ($E_R$) without developing symptoms and then become permanently immune to the virus ($R$).

$$\frac{dS^p}{dt} = -\beta^p\frac{S^pI^p}{N^p} - \kappa S^p$$

$$\frac{dE_R^p}{dt} = (1-\rho)\beta^p\frac{S^pI^p}{N^p} - (\varepsilon_R + \kappa)E_R^p$$

$$\frac{dE_I^p}{dt} = \rho\beta^p\frac{S^pI^p}{N^p} - (\varepsilon_I + \kappa)E_I^p$$

$$\frac{dI^p}{dt} = \varepsilon_I E_I^p - (\mu + \kappa)I^p$$

$\frac{dR^p}{dt} = \varepsilon_R E_R^p - \kappa R^p,$

where $\kappa = d + (b-d)\frac{N}{K}$ is the density dependent death rate, $N$ is the total bats population and $K$ is the carrying capacity. In Avenc Davì, Castanya, Summer refuges and Other caves the birth rate is $b = 0$ so $\kappa = d$. In Can Palomeres we model the birth rate pulse $b$, so $\kappa > d$. As above, in Can Palomeres, the force of infection also considers the cross-species contribution. The intrinsic $R_0^p$ value, the basic reproductive number in the absence of migration per patch $p$, is obtained using the next generation matrix approach:

$$R_0^p = \frac{\rho\beta^p\varepsilon_I}{(\varepsilon_I+\kappa)(\mu+\kappa)} \tag{3}$$

The force of infection for intra-species transmission in a given patch $p$ at time $t$ is:

$$\lambda^p(t) = R_0^p \frac{(\varepsilon_I+\kappa)(\mu+\kappa)}{\rho\varepsilon_I}\frac{I^p(t)}{N^p(t)} \tag{4}$$

As above, in Can Palomeres, the force of infection also considers the cross-species contribution, $R_0^{mix}\frac{(\varepsilon_I+\kappa)(\mu+\kappa)}{\rho\varepsilon_I}\frac{I_{spe_2}^{CP}(t)}{N_{spe_2}^{CP}(t)}$, where $R_0^{mix} = \alpha R_0^{CP}$.

**2.3 Model 3**

The transmission dynamics of model 3 is equivalent to the one described for model 2 with the addition of temporary immunity of average duration $\omega^{-1}$:

$$\frac{dS^p}{dt} = -\beta^p\frac{S^pI^p}{N^p} - \kappa S^p + \omega R^p$$

$$\frac{dE_R^p}{dt} = (1-\rho)\beta^p\frac{S^pI^p}{N^p} - (\varepsilon_R+\kappa)E_R^p$$

$$\frac{dE_I^p}{dt} = \rho\beta^p\frac{S^pI^p}{N^p} - (\varepsilon_I+\kappa)E_I^p$$

$$\frac{dI^p}{dt} = \varepsilon_I E_I^p - (\mu+\kappa)I^p$$

$$\frac{dR^p}{dt} = \varepsilon_R E_R^p - \kappa R^p + \omega R^p$$

Equations for Can Palomeres have in addition the birth rate pulse $b$ and the cross species interaction, as for model 2. The reproductive number per patch $R_0^p$ in absence of migration is equal to Eq. (3).

#### 2.4 Parameter values

We provide here a table listing the model parameters and their values.

**Table S5. Parameters description and values.**

| | **Parameter description** | ***M. myotis*** | | ***M. schreibersii*** | |
|---|---|---|---|---|---|
| | | *Default value* | *Ref.* | *Default value* | *Range* |
| $N$ | Total population size | 500 [400-600] | * | 17,000* | [16,000-18,000] |
| $\varepsilon_I^{-1}$ | Average incubation period leading to infectiousness | 30 days | [9] | 30 days | |
| $\varepsilon_R^{-1}$ | Average incubation period leading to immunity (models 2, 3) | 15 days | [10] | 15 days | |
| $\mu^{-1}$ | Average infectious period | 5 days | [3, 9] | 5 days | 2.5, 10 days |
| $\omega^{-1}$ | Average immunity period (models 1, 3) | 2 years | [9] | – | [180 – 730] days |
| $\rho$ | Proportion of exposed becoming infectious (models 2, 3) | 0.15,0.35, 0.5 | [10] | 0.15, 0.35, 0.5 | |
| $d$ | Natural death rate | 1/15 years$^{-1}$ | as for *M. schreibersii* | 1/15 years$^{-1}$ | [11] |
| $b$ | Pulsing birth rate (birthing season only) | $b(t) = d$ in birthing season $b(t) = 0$ otherwise | as for *M. schreibersii* | $b = d$ in Can Palomeres $b = 0$ in other caves | [11] |
| $\alpha$ | Degree of cross-species interaction | 0 (non-mixing), 0.5 (mixing); range: [0 – 1] | | | |

*From empirical estimates, see Table S1

### 3. Simulation details

#### 3.1 Stochastic and discrete integration of the disease dynamics

In each patch $p$ for a compartment $X^p$ we extract a random variable from a multinomial distribution for each possible transition out of the compartment in the discrete time interval $\Delta t$. Such variable, $\mathcal{D}^p(X^p, Y^p)$, determines the number of transitions from the compartment $X^p$ to a given compartment $Y^p$ occurring in $\Delta t$. The change of population size of a compartment in the time interval because of disease dynamics is given by the sum over all random variables extracted, each for a specific transition:

$$\Delta X^p = \sum_Y [-\mathcal{D}^p(X^p, Y^p) + \mathcal{D}^p(Y^p, X^p)] \quad (5)$$

Here we consider the evolution of the susceptible compartment $S^p$ of model 1 as a concrete example. All possible transitions from this compartment are: to the exposed $E_I^p$ and to the natural death given by the demography.

The random variables for these transitions from $S^p$ are extracted from the multinomial distribution:

$$Pr^{multin}\left(S^p, P_{S^p \to E_I^p}, P_{S^p \to death}\right) \quad (6)$$

with the transition probabilities:

- $P_{S^p \to E_I^p} = \lambda \Delta t$

- $P_{S^p \to death.} = -d\Delta t$

These three transitions cause a reduction of the size of that compartment $S^p$. The increase is given by the transition from the recovered $R^p$ to the susceptible $S^p$ after the loss of immunity (here we do not consider Can Palomeres where an additional transition refers to the birth rate). This transition is modeled by a random number extracted from a binomial distribution:

$$Pr^{bin}(R^p, P_{R^p \to S^p}) \quad (7)$$

with probability $P_{R^p \to S^p} = \omega\Delta t$, and $R^p$ being the number of recovered at time $t$ in patch *p*. After extracting these numbers from the corresponding distributions, we can calculate the stochastic variation of the population size of compartment $S^p$:

$$\Delta S^p(t) = S^p(t+1) - S^p(t) = -\left[\mathcal{D}^p\left(S^p, E_I^p\right) + \mathcal{D}^p(S^p, death)\right] + \mathcal{D}^p(R^p, S^p) \quad (8)$$

$\Delta t$ is equal to 1 day.

#### 3.2 Stochastic migration of *M. schreibersii*

The yearly migration of *M. schreibersii* is defined by seasonal movements (Table S2) with an integration time scale of 1 day. The number of *M. schreibersii* in the compartment $X^p$ traveling from patch $p$ to patch $p'$ in the discrete time interval $\Delta t$ is an integer random variable extracted from a multinomial distribution. As a concrete example let us consider the spring migration out of Avenc Davì of bats in the $X^{AD}$ compartment. The two possible destinations are the compartment $X^C$ in Castanya and the compartment $X^{OC}$ in Other caves. The random variables are extracted from:

$$Pr^{multin}\left(X^{AD}, P_{X^{AD} \to X^C}, P_{X^{AD} \to X^{OC}}\right) \quad (9)$$

with migration probabilities:

- $P_{X^{AD} \to X^C} = \eta^{AD \to C}\phi^{AD \to C}\Delta t$
- $P_{X^{AD} \to X^{OC}} = (1 - \eta^{AD \to C})\phi^{AD \to OC}\Delta t$

where: $X^{AD}$ is the number of *M. schreibersii* in Avenc Davì in the $X$ compartment at time *t*; $\phi^{AD \to OC}$ and $\phi^{AD \to C}$ are the daily migration rates from Avenc Davì to Castanya and from Avenc Davì to Other caves, respectively (Table S2); $\eta^{AD \to C}$ and $(1 - \eta^{AD \to C})$ are the estimated percentages of bats that migrate to the two destinations, respectively (Table S2). After extracting these numbers from the corresponding distributions, we calculate the change in the $X^{AD}$ population.

### 4. Maximum Likelihood

Likelihood was computed by assuming a binomial probability of testing positive, namely
$P(n_{p,t}|\omega, R_0) \sim Bin\left(N_{p,t}, r_{p,t}(\omega, R_0)\right)$,
where $N_{p,t}$ is the number of samples collected in the cave $p$, at time $t$; $n_{p,t}$ the ones tested positive; and $r_{p,t}(\omega, R_0)$ the average density of recovered predicted by the model. $n_{p,t}$ are considered to be statistically independent.

### 5. Additional results

In this section we provide additional numerical results mentioned in the main paper.

**Table S6. EBLV-1 persistence probability for *Miniopterus schreibersii* and *Myotis myotis* with the alternatives infection dynamics and transmission types considered.**

| Model | Transmission | $\rho$ | EBLV-1 persistence probability in *M. schreibersii* | | EBLV-1 persistence probability in *M. myotis* | |
|---|---|---|---|---|---|---|
| | | | Non-mixing | Mixing | Non-mixing | Mixing |
| Model 2 | frequency dependent | 0.15 | 0 | 0 | 0 | 0 |
| | | 0.35 | 0 | 0 | 0 | 0 |
| | | 0.5 | 0 | 0 | 0 | 0 |
| Model 3 | frequency dependent | 0.15 | 0 | $< 0.09$ | 0 | $< 0.09$ |
| | | 0.35 | 0 | $< 0.06$ | 0 | $< 0.04$ |
| | | 0.5 | 0 | 0 | 0 | 0 |
| Model 1 | density dependent | - | 0 | 0 | 0 | 0 |
| Model 2 | density dependent | 0.15 | 0 | 0 | 0 | 0 |
| | | 0.35 | 0 | 0 | 0 | 0 |
| | | 0.5 | 0 | 0 | 0 | 0 |
| Model 3 | density dependent | 0.15 | 0 | 0 | 0 | 0 |
| | | 0.35 | 0 | 0 | 0 | 0 |
| | | 0.5 | 0 | 0 | 0 | 0 |

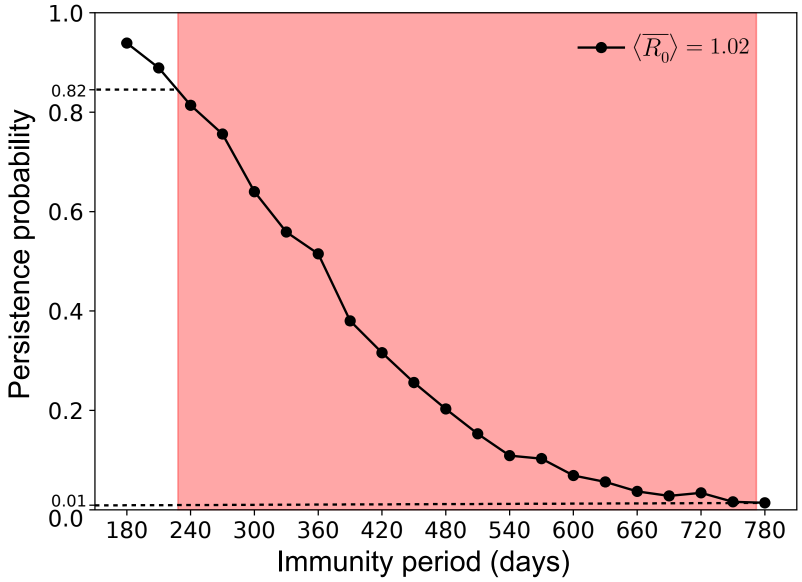

**Figure S1. EBLV-1 Persistence probability in the maximum likelihood estimate point.** Persistence probability of EBLV-1 in *M. schreibersii* population as a function of the immunity period with $\langle \overline{R_0} \rangle$ set to its maximum likelihood estimate. The red shaded area represent the 95% CI of the maximum likelihood estimate for the waning of immunity. The horizontal dashed lines indicate the persistence probability in the *M. schreibersii* population at the two extremes of the 95% CI of the waning of immunity.

### 6. Sensitivity analysis

In this section we provide a set of numerical results where we evaluated the sensitivity of the model predictions to variations of model parameters or data estimates.

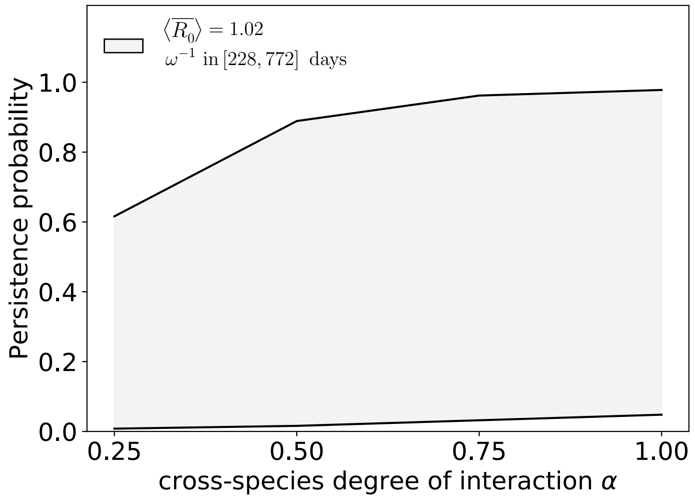

**Fig S2. Sensitivity analysis on the degree of cross-species mixing.** Persistence probability of EBLV-1 in *M. schreibersii* population as a function of the cross-species interaction $\alpha$ computed for the maximum likelihood estimate for $\langle \overline{R_0} \rangle$ and for values of the immunity period $\omega^{-1}$ spanning the estimated confidence interval.

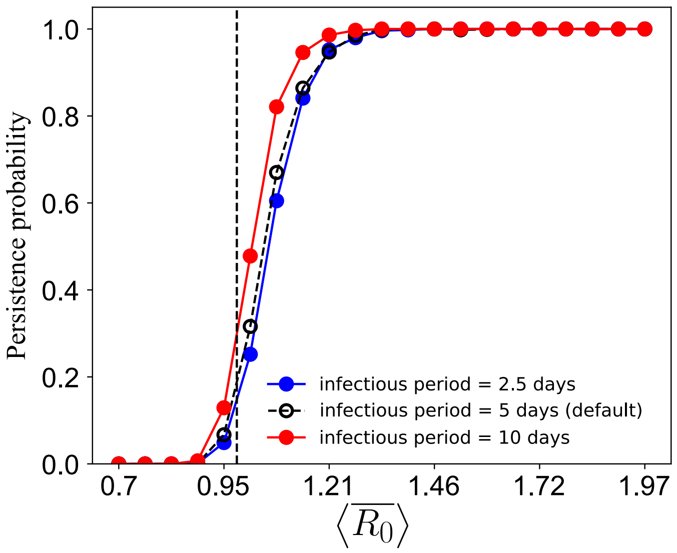

**Figure S3. Sensitivity analysis on the duration of the infectious period.** Persistence probability of EBLV-1 in *M. schreibersii* population for an infectious period of 2.5 days (blue) or of 10 days (red) compared to the predictions obtained with the default value of 5 days. The lines are computed on the maximum likelihood estimate for the waning of immunity. The vertical dashed line indicates $\langle \overline{R_0} \rangle = 1$.

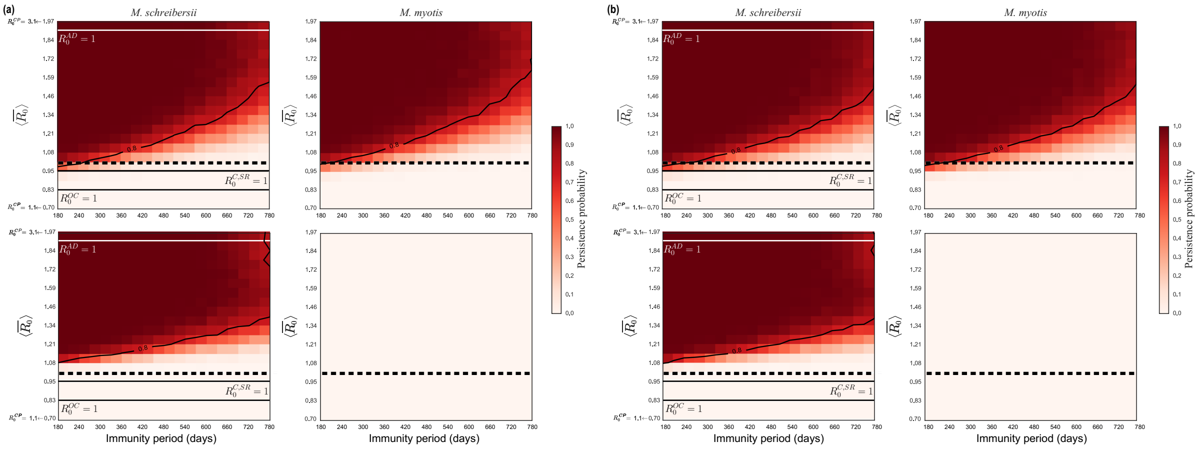

**Figure S4. Sensitivity analysis on *M. schreibersii* bats population size.** Persistence probability of EBLV-1 in *M. schreibersii* and in *M. myotis* as a function of the average reproductive number of the metapopulation system $\langle \overline{R_0} \rangle$ and of the immunity period $\omega^{-1}$, in mixing (top) and non-mixing (bottom) conditions, for: (a) a population of 16,000 *M. schreibersii* and 500 *M. myotis*; (b) a population of 18,000 *M. schreibersii* and 500 *M. myotis*. Contour lines indicate a persistence probability of 80%. The dashed horizontal line refers to $\langle \overline{R_0} \rangle = 1$. Solid horizontal lines refer to threshold conditions ($R_0^p = 1$) for the all caves.

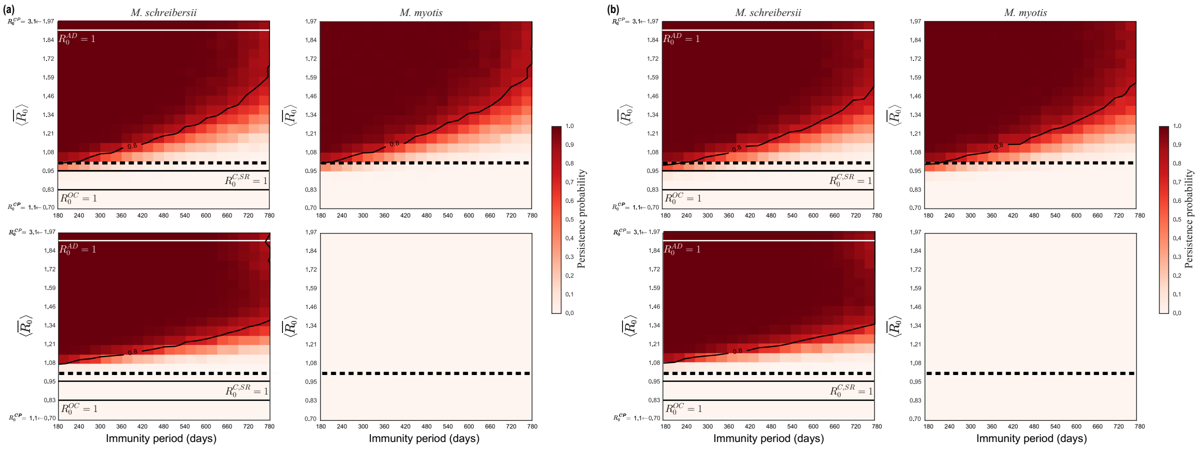

**Figure S5. Sensitivity analysis on *M. myotis* bats population size.** Persistence probability of EBLV-1 in *M. schreibersii* and in *M. myotis* as a function of the average reproductive number of the metapopulation system $\langle \overline{R_0} \rangle$ and of the immunity period $\omega^{-1}$, in mixing (top) and non-mixing (bottom) conditions, for: (a) a population of 17,000 *M. schreibersii* and 400 *M. myotis*; (b) a population of 17,000 *M. schreibersii* and 600 *M. myotis*. Contour lines indicate a persistence probability of 80%. The dashed horizontal line refers to $\langle \overline{R_0} \rangle = 1$. Solid horizontal lines refer to threshold conditions ($R_0^p = 1$) for the all caves.

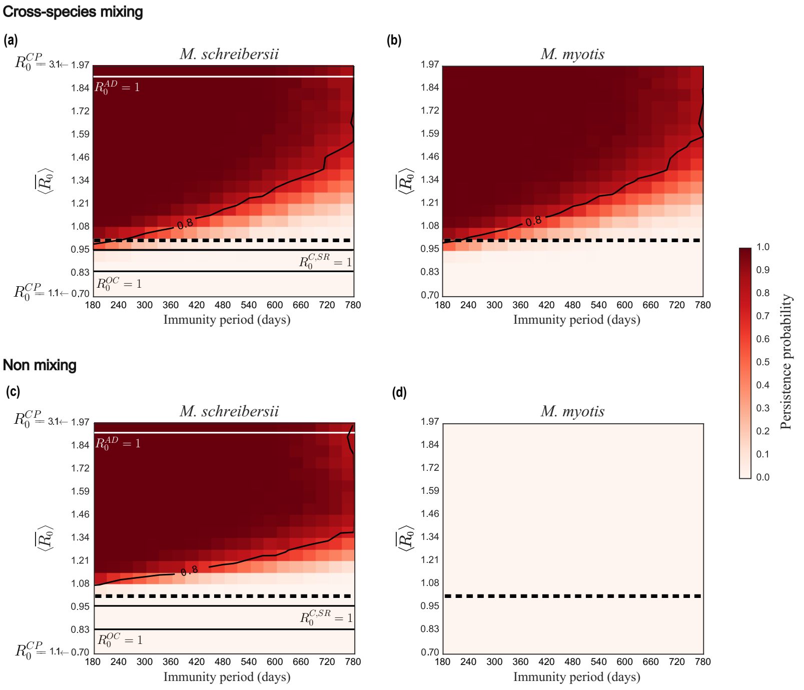

**Figure S6. Sensitivity analysis on starting date of migration events.** Persistence probability as a function of the average reproductive number of the metapopulation system $\langle\overline{R_0}\rangle$ and of the immunity period $\omega^{-1}$ for *M. schreibersii* and for *M. myotis* in the mixing scenario (top) and for non-mixing conditions (bottom) for starting dates of migration changed by a factor $\epsilon$, where $\epsilon$ is randomly extracted from a Gaussian distribution with a zero mean and a standard deviation of 1 week. The dashed horizontal line refers to $\langle\overline{R_0}\rangle = 1$. Solid horizontal lines refer to threshold conditions ($R_0^p = 1$) for the all caves.

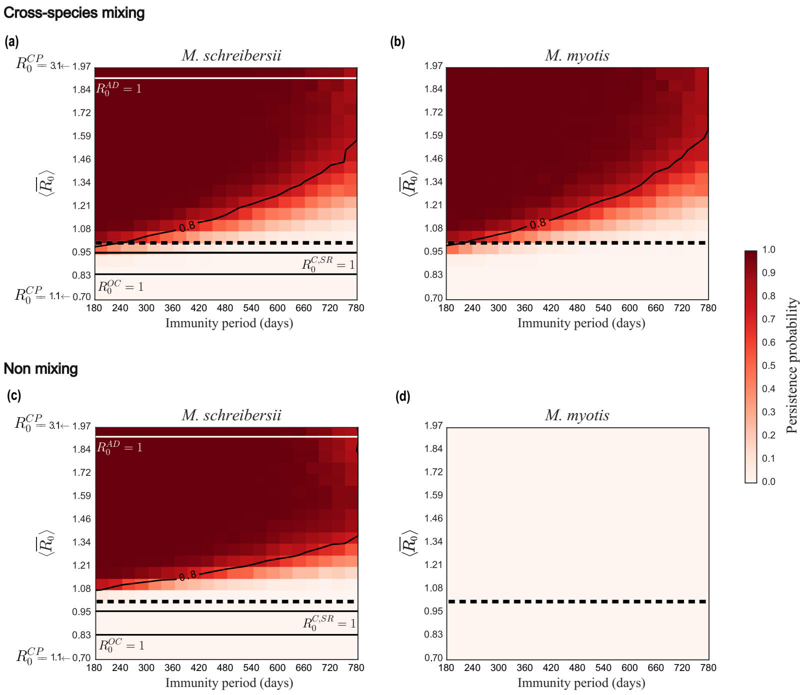

**Figure S7. Sensitivity analysis on duration of migration events.** Persistence probability as a function of the average reproductive number of the metapopulation system $\langle\overline{R_0}\rangle$ and of the immunity period $\omega^{-1}$ for *M. schreibersii* and for *M. myotis* in the mixing scenario (top) and for non-mixing conditions (bottom) for migration durations $\Delta t$ changed by a factor $\epsilon$, where $\epsilon$ is randomly extracted from a Gaussian distribution with a zero mean and a standard deviation of 1 week. The dashed horizontal line refers to $\langle\overline{R_0}\rangle = 1$. Solid horizontal lines refer to threshold conditions ($R_0^p = 1$) for the all caves.

11. J Serra-Cobo. Biological and ecological study of the Miniopterus schreibersii. [PhD thesis]. 1989.
